# Diversification of budding yeast polarization dynamics after compensatory experimental evolution

**DOI:** 10.64898/2026.11.04.764177

**Authors:** Marieke M. Glazenburg, Liedewij Laan

## Abstract

Essential cellular processes evolve under strong selection, yet protein networks underlying conserved functions can vary substantially. To what extent evolution constrains the intracellular dynamics of proteins performing essential functions is currently largely unknown. Here, we characterize budding yeast polarization phenotypes along a previously established evolutionary recovery trajectory, compensating for the loss of near-essential polarity gene *BEM1*. Through quantitative live cell imaging, we show that loss of *BEM1* weakens and destabilizes polarization. Surprisingly, in recovered evolved lineages, protein dynamics change significantly compared to the ancestor: most notably, presence of main regulator Cdc42-GTP is strongly decreased, while polarisome member Spa2 localizes more slowly into mobile foci. Although separate phenotypic characteristics are only weakly constrained, we find evidence of effective protein-independent localization modes, revealing signs of convergence along the evolutionary trajectory. Together, our results show that evolution can restore global cellular function without conserving microscopic organization, instead giving rise to novel molecular phenotypes.

## INTRODUCTION

Many intracellular processes are governed by complex, self-organizing networks of highly interactive proteins^1–3^. Although the functionality of such processes is often conserved across evolution, the underlying molecular networks can vary substantially between species and over evolutionary time^4–6^. This is true even for essential functions, such as chromosome segregation, endocytosis or certain metabolic pathways, which evolve under strong selection pressure^7–10^. Thus, while core cellular functions are robustly maintained across evolution, the protein networks orchestrating these functions remain adaptable.

While genetic diversity underlying cellular functions has been mapped out in great detail, much less is known about the variability of intracellular protein dynamics across evolution. Protein localization patterns are strong determinants of cellular function^11–13^, suggesting that their dynamics may be evolutionarily constrained. At the same time, molecular localization is governed by complex interactions between the proteins and their physio-chemical environment^14–16^, making the relation between selectable mutations and molecular phenotype non-trivial. How evolution shapes protein dynamics under functional and physio-chemical constraints is currently an open question. Recent work on fungal endocytosis has demonstrated how even conserved molecular components display distinct dynamics across different species, varying in arrival order, timescales and membrane remodeling characteristics^10^, suggesting that specific protein behavior can change significantly depending on the genetic background while the core cellular function is conserved. How these observations extend to other cellular processes or to more subtle genetic variation within the same species is largely unknown.

Polarization in the budding yeast *Saccharomyces cerevisiae* is a prime example of an evolutionarily flexible cellular process and therefore a powerful model system in this context^17^. Polarization, which in budding yeast refers to the process of determining the site of daughter cell emergence, is required for asymmetric cell division and thus comprises an essential step in the budding yeast life cycle^18–20^. Despite this essential role, the composition of fungal polarity networks is highly variable across natural evolution^6,17^, suggesting that polarization as a cellular function is conserved without fixing specific molecular details.

Besides natural evolution, the evolutionary flexibility of yeast polarization has also been demonstrated in laboratory settings. It has previously been shown that budding yeast is capable of recovering from near-fatal gene losses, including polarity genes, through experimental evolution^21^. In the context of polarity specifically, cells are able to recover from the loss of near-essential scaffold gene *BEM1* by a reproducible trajectory of subsequent loss-of-function mutations in *BEM3* and *NRP1*^22^. Remarkably, a recent genome-wide mutation screen demonstrated that almost 900 genes (approximately 13% of the total number of annotated genes in the genome) undergo a significant change in disruption tolerance across this evolutionary trajectory, implying a global rewiring of the polarization network^23^. These observations raise the question: when evolution recovers a cellular function through extensive genetic rewiring, are the underlying protein dynamics also restored to their ancestral state, thereby constraining evolution to particular molecular phenotypes, or can the same cellular function be repaired by adopting novel molecular behavior?

Here, we aim to address this question using the evolutionary recovery of cell polarization following the deletion of *BEM1* as a model. Using quantitative live cell imaging^24^, we characterize polarization phenotypes along the evolutionary trajectory, comparing protein dynamics in the perturbed Bem1-deficient state to several evolved recovered lineages that have regained near-ancestral growth characteristics and morphology. Strikingly, the localization patterns of polarizing proteins in evolved cells are not reverted to the ancestral phenotype. Instead, evolved mutants reproducibly exhibit different dynamics that are most similar to the perturbed state prior to evolutionary repair, suggesting that even closely related mutants can employ highly distinct but similarly efficient molecular strategies for polarization. Moreover, by describing polarization phenotypes in terms of quantitative characteristics, we identify effective protein localization ‘modes’ corresponding to different strategies of signal accumulation during polarization. The trajectories of these localization modes converge across evolutionary recovery, suggesting that evolution responds to deficiencies in protein recruitment efficiency by balancing total localization timescales. Our findings demonstrate how evolution can functionally conserve cellular processes without maintaining specific dynamic behavior of underlying components.

## RESULTS

### Population-level characteristics are rapidly recovered

We commenced by assessing to what extent the population as a whole restores fitness and cell morphology over the course of evolutionary recovery, reflecting how efficiently the cellular function of polarization is repaired. We focused on the reproducible recovery following *BEM1* deletion using the previously characterized *bem1*Δ *bem3*Δ *nrp1*Δ mutant^22,23^, from hereon referred to as the reconstructed evolved mutant (REM). Additionally, we studied three independently evolved lineages from the original evolution experiment^22^, containing loss-of-function mutations in *BEM3* and other compensatory mutations but not in *NRP1*, from hereon referred to as EV1-3 (Fig. 1(a)). In line with what has been previously shown^22^, population growth was significantly delayed in *bem1*Δ mutants (Fig. 1(b) and Fig. S1). This defect was largely alleviated by additional knockout of *BEM3* and further recovered by deletion of *NRP1*, resulting in a doubling time in the final reconstructed evolved *bem1*Δ *bem3*Δ *nrp1*Δ mutant that was only roughly 10% elevated compared to the ancestor (Fig. 1(b)). Likewise, evolved mutants EV1-3 had doubling times in a similar range (Fig. 1(b)). Hence, defects in proliferation of the population are largely recovered at the end of the evolutionary trajectory.

**Figure 1:**
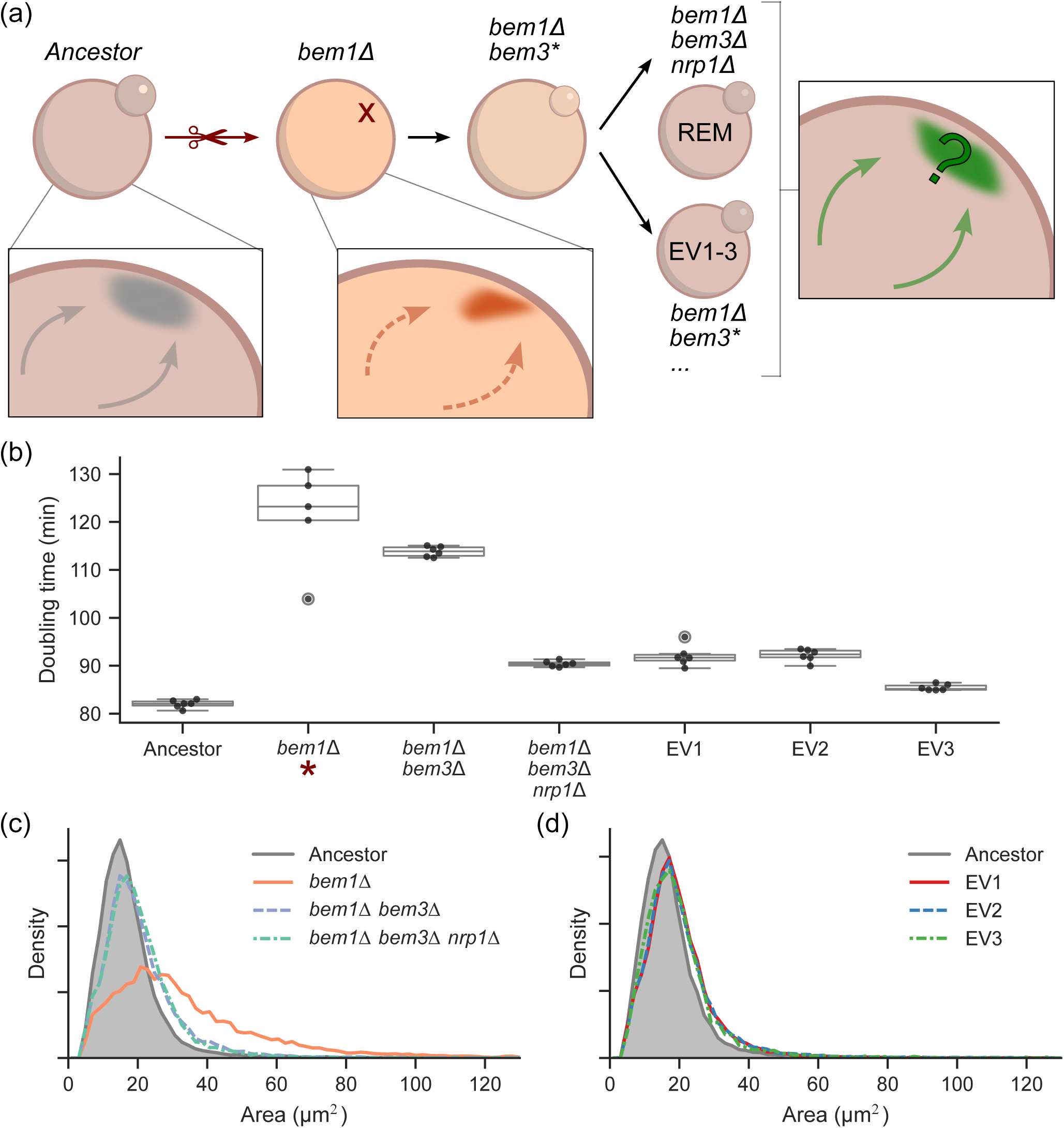
Evolutionary recovery of population-wide characteristics. (a) Illustration of the evolutionary trajectory, starting with knock-out of *BEM1*, which perturbs polarization and budding, and recovering by inactivation of *BEM3* and subsequently *NRP1* (in the reconstructed evolved mutant, REM) or other mutations (in evolved lines EV1-3). Functionality of polarization and budding is restored in the evolved cells, but protein localization dynamics are unknown. (b) Log phase doubling times of the ancestor, the perturbed *bem1*Δ mutant and recovered mutants. All strains express Spa2-Citrine. Note that the derived doubling time of *bem1*Δ cells (marked with a red star) is underestimated in bulk growth assays, since the stage of fast logarithmic growth used for fitting doubling times is readily dominated by cells with suppressor mutations; significant growth delay is still visible in the increased lag time (Fig. S1). (c) Distributions of cell sizes along the reconstructed recovery trajectory, compared to the ancestor (grey). All strains express Spa2-Citrine. (d) Distributions of cell sizes of evolved mutants EV1-3 expressing Spa2-Citrine, compared to the ancestor (grey).

Next, we expanded upon the growth rate analysis by asking whether evolutionary recovery has a discernible effect on cellular morphology, since aberrant morphology can be an indicator of polarization defects^25,26^. To address this question, we performed a morphology screen using brightfield images of over 2000 cells per condition. As shown in Fig. 1(c), the *bem1*Δ population had an increased fraction of larger cells, likely corresponding to cells that fail to divide and continue growing isotropically until lysis. However, this phenotype disappeared following deletion of *BEM3*, which recovered near-ancestral cell sizes. Additional deletion of *NRP1* had no discernible effect, leaving a small overall increase in cell size compared to the ancestor. Similarly, evolved mutants EV1-3 also exhibited close to ancestral cell sizes (Fig. 1(d)). Additionally, cells were only minimally elongated in both the initial *bem1*Δ state and in all evolved mutants (Fig. S2). Therefore, we conclude that defects in the population-level phenotype are rapidly recovered over evolutionary repair, thus restoring the global functionality of polarization to near-ancestral levels.

### Bem1-deficient cells can break symmetry but polarization is unstable

We have seen that deletion of *BEM1* severely disrupts cellular proliferation, but is not fully lethal. To understand what defects occur in polarization in the absence of Bem1, we characterized polarization dynamics using two well-studied polarity proteins: Cdc42, a small highly conserved RhoGTPase that is considered the main regulator of polarity in yeast^27^, and Spa2, a scaffold that is part of the polarisome directing membrane growth towards the polarized zone^28,29^. Cdc42 is localized through multiple positive feedback loops, one of which is mediated by Bem1^30–34^. Downstream effectors of GTP-bound Cdc42 promote localization of Spa2, a scaffold component of the polarisome driving nucleation of polarized actin cables, which in turn mediate localization of Cdc42 and delivery of vesicles towards the polarized zone^29,35–37^. Thus, Cdc42 and Spa2 can serve as relevant representatives of both signaling and cytoskeletal organization during polarization.

We imaged Spa2-Citrine in *bem1*Δ cells and compared this to previously obtained data of Cdc42-sfGFP in Bem1-depleted cells^24^ (referred to as *Bem1-*), in which Bem1 was degraded using an auxin inducible degron. Imaging revealed that the majority of Bem1-deficient cells was able to break symmetry and polarize both Cdc42 and Spa2. However, defects occurred in regulating the maintenance of the polarity site. Cells exhibited diverse polarization phenotypes, including normal polarization with successful budding, continuous polarization in elongated cells, prematurely dissociating spots without bud growth and occasional failure to polarize at all (Fig. 2(a)). These phenotypes were present in both Cdc42 (as previously demonstrated in^24^) and Spa2, while Spa2 also occasionally formed persistent non bud-inducing spots (Fig. S3(a)). Thus, although Bem1-deficient cells can induce polarization, there are defects in the regulation of polarity maintenance, either resulting in premature depolarization or a failure to depolarize when necessary.

**Figure 2:**
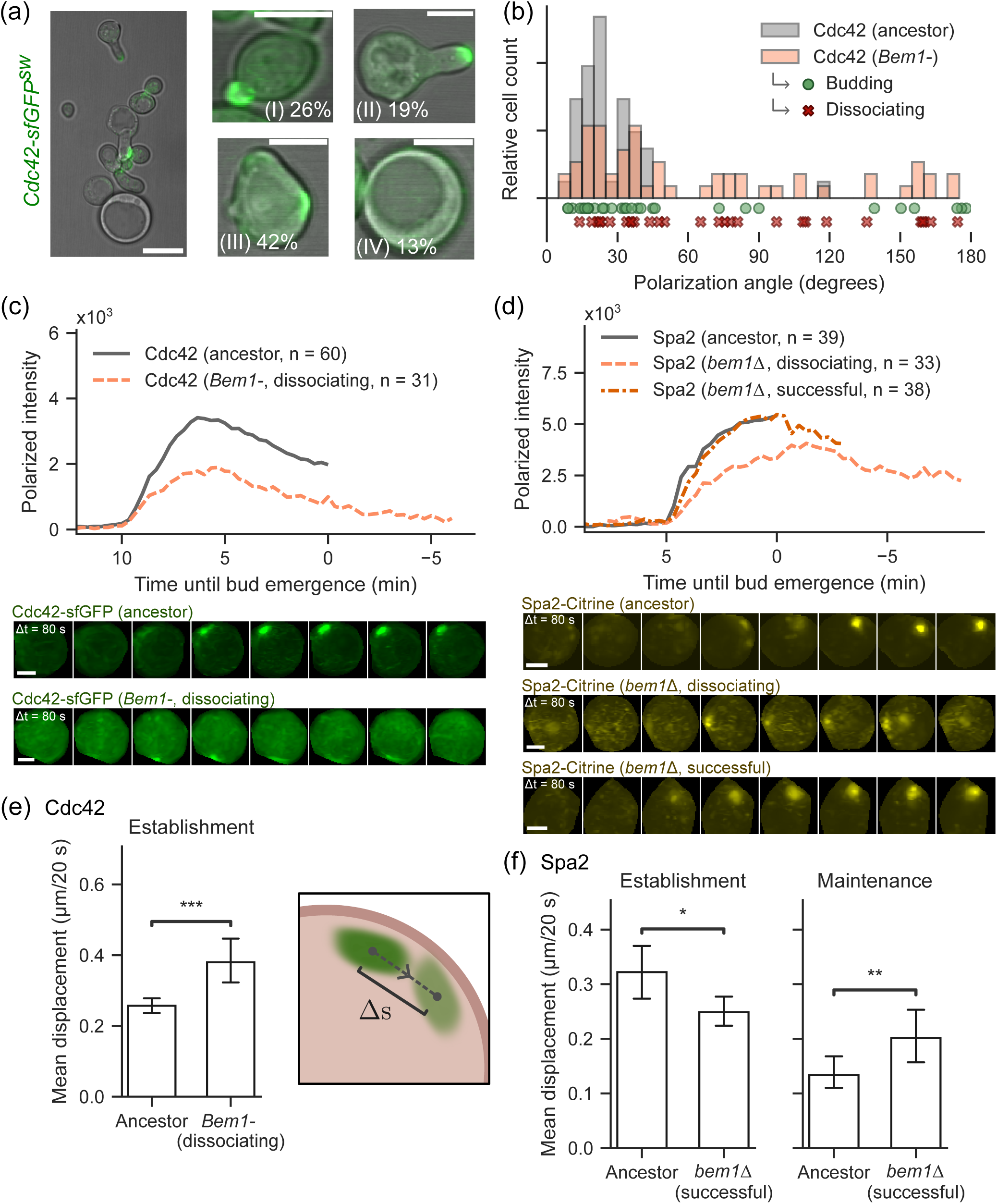
Polarization phenotype of Bem1-deficient cells. (a) Representative maximum projected confocal micrographs of Bem1-depleted (*Bem1-*) cells expressing Cdc42-sfGFP^sw^ (green) overlaid with brightfield images. Left: large field of view (scale bar 10 µm). Right: examples of polarization phenotypes (scale bars 5 µm). (I) Normal polarization and growth, (II), polarization resulting in elongated cell growth, (III) dissociating polarity spots and (IV) large unpolarized cells. Percentages reflect phenotype occurrence in a random count of n = 117 cells. Adapted from^24^. (b) Polarization angles with respect to the previous division site of Cdc42-sfGFP^sw^ in the wildtype ancestor (grey) and Bem1-depleted cells (*Bem1-*, orange). Circles and crosses indicate whether polarization events in Bem1-depleted cells initiated successful daughter cell growth. (c) Mean polarized intensity trace of Cdc42-sfGFP^sw^ in the ancestor (grey, n = 60) and dissociating spots in Bem1-depleted cells (orange, n = 31), aligned at the onset of polarization. Adapted from^24^. Individual traces are shown in Fig. S4. Representative micrographs of single polarizing cells are shown below (maximum projections, scale bars 2 µm). (d) Mean polarized intensity trace of Spa2-Citrine in the ancestor (grey, n = 39) and dissociating (orange, n = 33) and successful spots (red, n = 38) in *bem1*Δ cells, aligned at the onset of polarization. Time axis is relative to bud emergence in the ancestor. Individual traces are shown in Fig. S4. Representative micrographs of single polarizing cells are shown below (maximum projections, scale bars 2 µm). (e) Cdc42-sfGFP^sw^ spot mobility during polarity establishment expressed as mean displacement per 20 s time frame, in the ancestor and for dissociating spots in Bem1deficient cells. (f) Spa2-Citrine spot mobility during polarity establishment and maintenance, in the ancestor and for successful bud-inducing spots in *bem1*Δ cells. Error bars indicate 95% confidence intervals. Stars indicate significance in a Mann-Whitney U-test (\**p <* 0.05; \*\**p <* 0.01; \*\*\**p <* 0.001; \*\*\*\**p <* 0.0001; ns, not significant).

Additionally, Bem1-deficient cells showed defects in the relative positioning of the polarized spot. We confirmed that polarization of Cdc42 in Bem1-deficient cells takes place scattered across the membrane (Fig. 2(b)), consistent with the budding patterns observed in earlier work^38^. The same held true for Spa2 (Fig. S3(b)). Moreover, based on the positioning of successful and dissociating polarity spots, successful bud emergence could occur anywhere on the membrane, but was more likely in closer proximity to the bud scar (Fig. 2(b) and Fig. S3(b)): for Cdc42 in Bem1-deficient cells, 62% of events occurring at angles below 60*^◦^* led to successful budding (n = 34), compared to 38% of events at higher angles (n = 24).

Quantitative image analysis revealed that the defective polarization phenotypes can be linked to a reduction in polarization strength and stability. As we have shown previously^24^, dissociating spots of Cdc42 in Bem1-depleted cells exhibit dynamics that were comparable in timescale to successful polarization in the ancestor, but spots remained weaker before disassembling (Fig. 2(c)). Hence, Cdc42 polarity is overall established within a similar time frame, but the strength of polarization was reduced (see also S5(a)). Moreover, the growing polarity spots are less stably positioned compared to the ancestor, reflected in larger movement of the spot on the membrane (Fig. 2(e)). We then extended our analysis to Spa2, revealing that Spa2 also exhibited weaker localization in dissociating spots, although not as strongly reduced as for Cdc42 (Fig. 2(d)). Spots leading to successful budding on the other hand were similarly strong or even more prominent than the ancestral spots, but their total localization time was increased (Fig. 2(d) and Fig. S5(b)). Such successful spots also had an increased mobility during polarity maintenance (Fig. 2(f)), showing that wandering of the polarity site by itself does not necessarily prevent bud formation.

In summary, polarization in Bem1-deficient cells has decreased affinity to the bud scar and is characterized by dissociating spots that are weaker and more mobile. Hence, the ability to break symmetry by either of the two imaged polarity components is not fully lost in the perturbed state, but instead dynamics are qualitatively and quantitatively altered.

### Evolutionary recovery restores functionality but not dynamics of polarization

Having established how polarization is affected in the absence of Bem1, we turned our attention to the recovery of protein dynamics in the reconstructed evolved *bem1*Δ *bem3*Δ *nrp1*Δ mutant (REM). Since the Cdc42-sfGFP^sw^ allele causes growth rate defects in this background (Fig. S6), we instead made use of a well-established reporter for Cdc42-GTP^39,40^, the Gic2 PBD domain. First examining the site selection pattern of the REM, we observed that affinity to the bud scar is regained compared to the Bem1-deficient state: polarization was biased to the regime in closest proximity to the landmark, similar to the ancestor (Fig. 3(a) and Fig. S3(c)). However, although the frequency was reduced, the REM did maintain the propensity to polarize and bud further away from the previous division site, similar to the Bem1-deficient mutants. Thus, the tendency to break symmetry spontaneously was still present in the evolved state.

**Figure 3:**
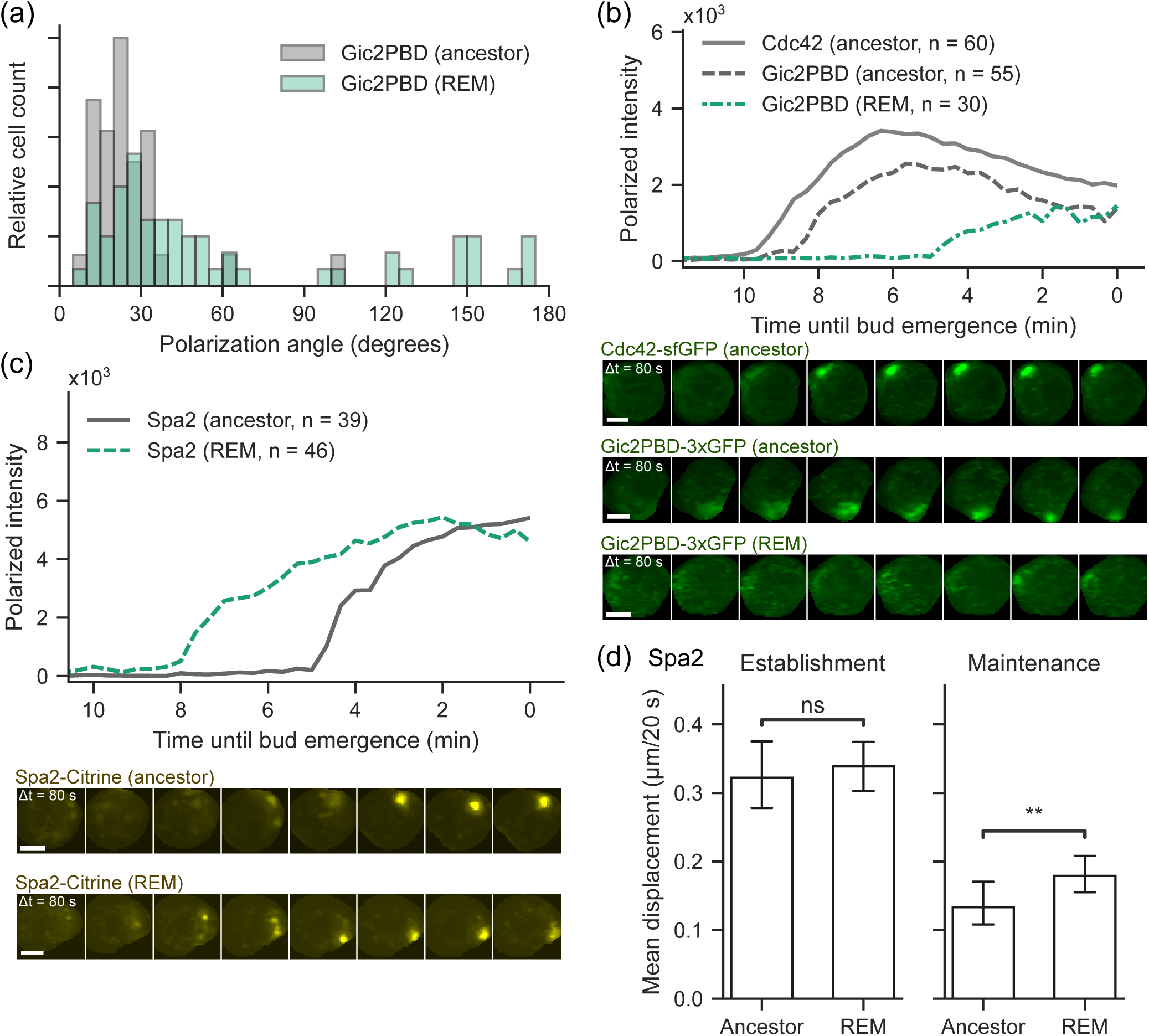
Polarization phenotype of the reconstructed evolved mutant. (a) Polarization angles with respect to the previous division site of Gic2PBD-3xGFP in the wildtype ancestor (grey) and reconstructed evolved mutant (blue). (b) Mean polarized intensity trace of Cdc42-sfGFP^sw^ (grey solid, n = 60) and Gic2PBD-3xGFP (grey dashed, n = 55) in the ancestor and Gic2PBD-3xGFP in the reconstructed evolved mutant (blue, n = 30), aligned at the bud emergence at the end of the timelapse. Individual traces are shown in Fig. S4. Representative micrographs of single polarizing cells are shown below (maximum projections, scale bars 2 µm). (c) Mean polarized intensity trace of Spa2-Citrine in the ancestor (grey, n = 39) and the reconstructed evolved mutant (blue, n = 46), aligned at bud emergence at the end of the timelapse. Individual traces are shown in Fig. S4. Representative micrographs of single polarizing cells are shown below (maximum projections, scale bars 2 µm). (d) Spa2-Citrine spot mobility during polarity establishment and maintenance, in the ancestor and the reconstructed evolved mutant (REM). Error bars indicate 95% confidence intervals. Stars indicate significance in a Mann-Whitney U-test (\**p <* 0.05; \*\**p <* 0.01; \*\*\**p <* 0.001; \*\*\*\**p <* 0.0001; ns, not significant).

Strikingly, polarization dynamics in the REM changed drastically. Rather than localizing well before bud emergence, Gic2PBD arrived later and at strongly reduced peak intensity, while the characteristic depolarization phase during spot maintenance had mostly vanished (Fig. 3(b) and Fig. S5(c)). Thus, while uncompromised in growth rate (Fig. S7), the REM showed significantly weakened polar localization of Cdc42-GTP. Although Spa2 is generally considered a downstream component of Cdc42 polarization^41^, dynamics of Spa2 in the evolved mutant did not match the decrease in polarization strength of Gic2PBD. Instead, localization was unaffected in final intensity but recruitment occurred at a lower rate (Fig. 3(c) and Fig. S5(d)), similar to the dynamics of successful spots in the *bem1*Δ background (Fig. 2(d)). Moreover, Spa2 fronts remained more mobile during the maintenance of the site (Fig. 3(d)), again mirroring Bem1-deficient dynamics. Notably, upon comparison of Fig. 3(b) and (c), the relative dynamics of Spa2 and Gic2PBD appear to invert in the REM: whereas the ancestor shows significant localization of Gic2PBD before and during localization of Spa2, the evolved mutant is characterized by an extended phase of strong Spa2 localization with minimal presence of Gic2PBD. Simultaneous imaging of Cdc42-GTP and Spa2 confirmed this observation (Fig. S8).

Together, these observations demonstrate that the deviating polarization dynamics observed in Bem1-deficient cells are not restored to ancestral behavior in the reconstructed evolved mutant. Importantly, the presence of Cdc42-GTP in the polar zone is strongly diminished, similar to the reduced localization strength of Cdc42 in dissociating spots in the Bem1-depleted background. Meanwhile, dynamics of Spa2 are comparable to the behavior of bud-inducing spots in the *bem1*Δ background, although the larger *bem1*Δ spots are restored to the smaller volumes of the ancestor (Fig. S5(d)).

The drastic changes in the localization of Gic2PBD could have multiple causes. First, it is possible that localization strength is decreased in the mutant due to less efficient binding of the PBD domain to Cdc42-GTP, for instance due to increased competition for the p21 binding domain (PBD) that constitutes the biosensor, decreasing localization of the probe despite unchanged levels of Cdc42-GTP. Although we cannot reject this possibility with certainty, the Gic2PBD sensor has been used extensively in a wide variety of genetic backgrounds^42–47^, and so far no suspicions of altered binding affinity have been raised. Another possibility is that total expression levels of the probe (expressed under the native Gic2 promoter) are reduced in the reconstructed evolved mutant. However, cytoplasmic fluorescence intensities indicated similar abundances in both backgrounds (Fig. S9). If we dismiss both alternative possibilities, we must conclude that Cdc42-GTP itself has a decreased polarized presence in the reconstructed evolved mutant. Remarkably, this would imply that Spa2 localizes mostly independently of Cdc42-GTP.

### Independent evolutionary trajectories converge to similar polarization phenotypes

Polarization dynamics in the reconstructed evolved mutant are significantly altered. Consequently, we wondered if these alterations are specific to this particular mutant strain or if the observed changes form a general response to the loss of *BEM1*. Therefore, we turned to evolved mutants EV1-3, which are Spa2-Citrine expressing clones derived from independent replicates of the experimental evolution with confirmed *BEM3* early stop mutations, but different subsequent rescue mutations (see Fig. 1(a)). Given the fact that mutants EV1-3 represent independent genetic solutions to the loss of *BEM1*, we asked to what extent their polarization dynamics differ from the reconstructed evolved mutant.

Much like in the REM, polarization of Spa2-Citrine in EV1-3 was mostly directed towards the bud scar, but occasional events occurred elsewhere on the membrane (Fig. S3(d)), indicating an inherent tendency for spontaneous symmetry breaking in all evolved mutants. Quantitatively, polarization in EV1 and EV3 closely resembled both the REM and successful spots in *bem1*Δ cells, showing prolonged localization and overall increased presence at the polar zone (Fig. 4(a) and Fig. S5(e)). Meanwhile, Spa2 in EV2 had a shorter polarization time that is more akin to the ancestor, although reaching a lower final intensity (Fig. 4(a)). Additionally, the mobility of the polarized zone during maintenance was elevated in all evolved mutants, indicating a universal feature of Bem1-independent polarization (Fig. 4(b)). Together, these observations demonstrate that the recovery phenotype of Spa2 polarization reproducibly exhibits extended polarization at lower recruitment rates into more mobile spots, thereby suggesting a general pattern in response to the loss of Bem1.

**Figure 4:**
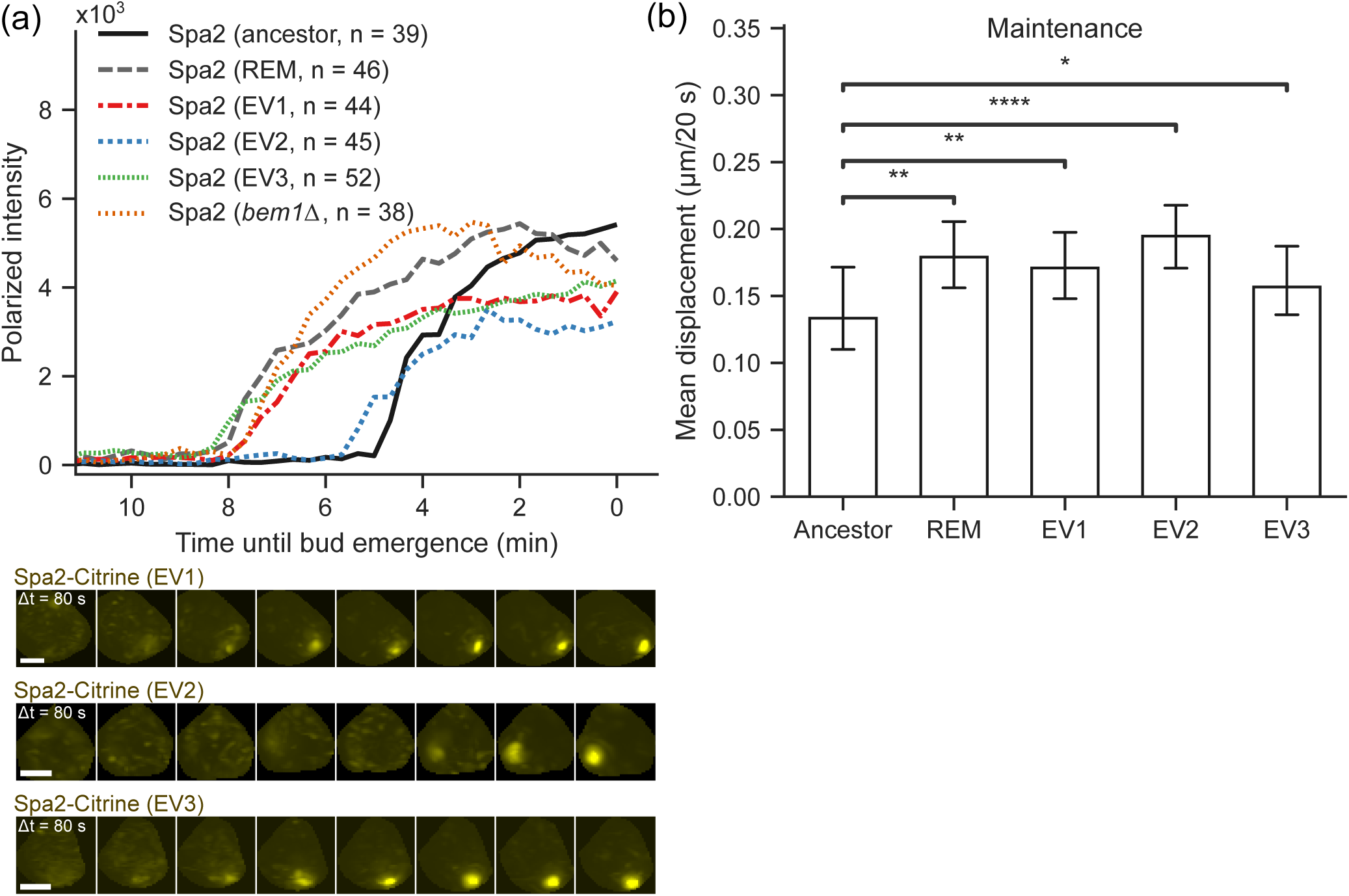
Polarization phenotypes of Spa2-Citrine in evolved mutants EV1-3. (a) Mean polarized intensity trace of Spa2-Citrine in the ancestor (solid grey, n = 39), the reconstructed evolved mutant (dashed grey, n = 46) and evolved mutants EV1 (red, n = 44), EV2 (blue, n = 45) and EV3 (green, n = 52), aligned at the bud emergence at the end of the timelapse. The mean trace of successful polarization in *bem1*Δ (orange, n = 38) is shown for reference. Individual traces are shown in Fig. S4. Representative micrographs of single polarizing cells are shown below (maximum projections, scale bars 2 µm). (b) Spa2-Citrine spot mobility during polarity maintenance, in the ancestor and the evolved lines EV1-3. Error bars indicate 95% confidence intervals. Stars indicate significance in a Mann-Whitney U-test (\**p <* 0.05; \*\**p <* 0.01; \*\*\**p <* 0.001; \*\*\*\**p <* 0.0001; ns, not significant).

### Evolutionary recovery trajectory converges on effective polarization modes

Evolutionary recovery following the loss of *BEM1* appears to affect polarization dynamics somewhat reproducibly, indicating potential constraints in polarization phenotypes. To investigate in more detail which aspects of polarization might be constrained, we make use of our previously developed framework^24^ describing polarization phenotypes by means of quantitative observables (illustrated in Fig. 5(a)). This way, each polarization event is represented by a set of observables, which include relevant timescales of polarity establishment and maintenance, recruitment rates and morphological characteristics of the polarized spot (see also Fig. S5). We then asked whether there are trends or constraints in the space spanned by these descriptive phenomenological observables.

**Figure 5:**
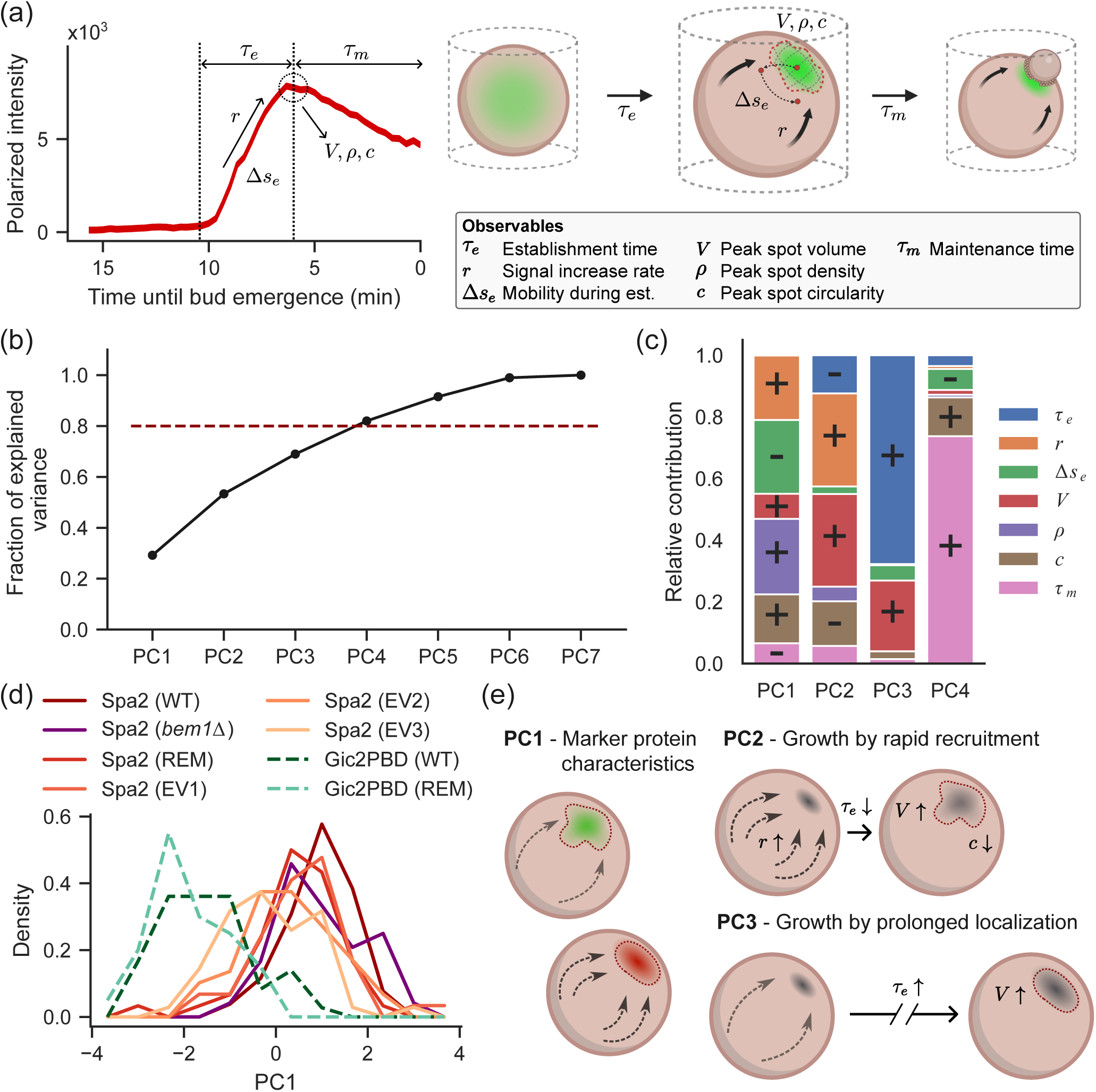
Identification of effective localization modes by PCA. (a) Summary of the quantitative characteristics from^24^ used to describe polarization phenotypes. (b) Cumulative ratio of explained variance per principal component. Horizontal dashed line indicates 80% explained variance. (c) Compositions in terms of polarization characteristics of the first four principal components resulting from PCA performed on all Gic2PBD and Spa2 data from successful polarization events pooled. + and - signs indicate the relative sign of each contribution. Note that relative signs are only meaningful within the same PC, since each PC can be arbitrarily multiplied by a factor -1. (d) Distributions of each marker-background combination along PC1. REM = reconstructed evolved mutant. (e) Proposed interpretation of PC1 (protein-specific characteristics), PC2 (growth by rapid recruitment with loss of circularity) and PC3 (growth by extending total localization time).

Pairwise correlations between observables were generally weak (Fig. S10(a) and (b)), suggesting that many direct characteristics of polarization can vary independently. Nevertheless, it is possible that evolution does not constrain these characteristics in isolation, but instead affects effective phenotypic classes comprised of combinations of observables. To investigate whether the protein localization phenotypes along the evolutionary trajectory contain such behavioral modes, we performed linear principal component analysis (PCA) on the pooled sets of observables derived from successful Gic2PBD and Spa2 polarization events across all studied conditions. Essentially, this indicates whether a dimensional reduction of the phenotypic phase space is possible through linear combinations of the original observables.

The first four principal components (PCs) together explained *>*80% of the total variance (Fig. 5(b)), suggesting that the phase space of polarization observables can be described by a lower-dimensional representation and thus may be subject to structural constraints. First, the most dominant axis (PC1, comprised of a complex mixture of characteristics as shown in Fig. 5(c)) separated dynamics of Gic2PBD and Spa2 independently of the genetic background and can thus be interpreted as a measure of marker identity (Fig. 5(d)). This way, the other derived phenotypic modes were effectively decoupled from protein-specific qualities. Second, the smallest considered axis (PC4) was mainly composed of the maintenance time, corresponding to the time between peak polarity establishment and bud emergence, together with a small contribution of spot circularity, and can therefore be viewed as a measure of maintenance persistence of regularly shaped spots (Fig. 5(c)). This axis thus primarily separated phenotypes based on the duration of extended localization after initial establishment and before daughter cell growth, ranging from the extremely short timescale of Gic2PBD in the reconstructed evolved mutant to the more prolonged localization of Spa2 in most evolved mutants (Fig. S10(c)). Strikingly, the spot maintenance time had very little contribution to the other three axes (Fig. 5(c)), highlighting this characteristic as an independently varying property of polarization.

We then focused on the remaining two axes (PC2 and PC3), which appeared to capture distinct modes of protein accumulation. One axis (PC2) was comprised of increasing spot volume and localization rate at the cost of establishment time (Fig. 5(c)), which we interpreted as a measure of irregular spot growth by means of rapid recruitment. Conversely, the other axis (PC3) consisted of increasing volume together with an increasing establishment timescale (Fig. 5(c)) and therefore reflected a measure of growth through prolonging the total accumulation time. Since these two axes had been largely decoupled from protein identity by PC1, they effectively represented protein-independent modes of localization during polarization (Fig. 5(e)).

Strikingly, over the course of evolutionary recovery, both Gic2PBD and Spa2 moved to a similar area of the phase space spanned by the two localization modes (Fig. 6(a)). More precisely, the contribution of rapid recruitment (PC2) was decreased, while growth by prolonged accumulation (PC3) was balanced to a moderate level. We speculate that the decrease in the rapid recruitment component is caused by the loss of positive feedback-inducing *BEM1* and Cdc42-cycling regulators (e.g. *BEM3*) in the evolved mutants, decreasing overall turnover and thus the accessible recruitment rate. To compensate for this, polarization can be rescued by extending the total duration of accumulation, although there appear to be constraints to this extension favoring moderate timescales. This is exemplified by the trajectory of Spa2 through the phase space, first increasing and subsequently decreasing the contribution of growth by prolonged recruitment when transitioning from the ancestor through *bem1*Δ to the recovered evolved states (Fig. 6(a), solid arrows). Meanwhile, polarization of Gic2PBD already occupied a moderate timescale regime in the ancestor and mainly lost rapid recruitment capacity over evolutionary recovery (Fig. 6(a), dashed arrow).

**Figure 6:**
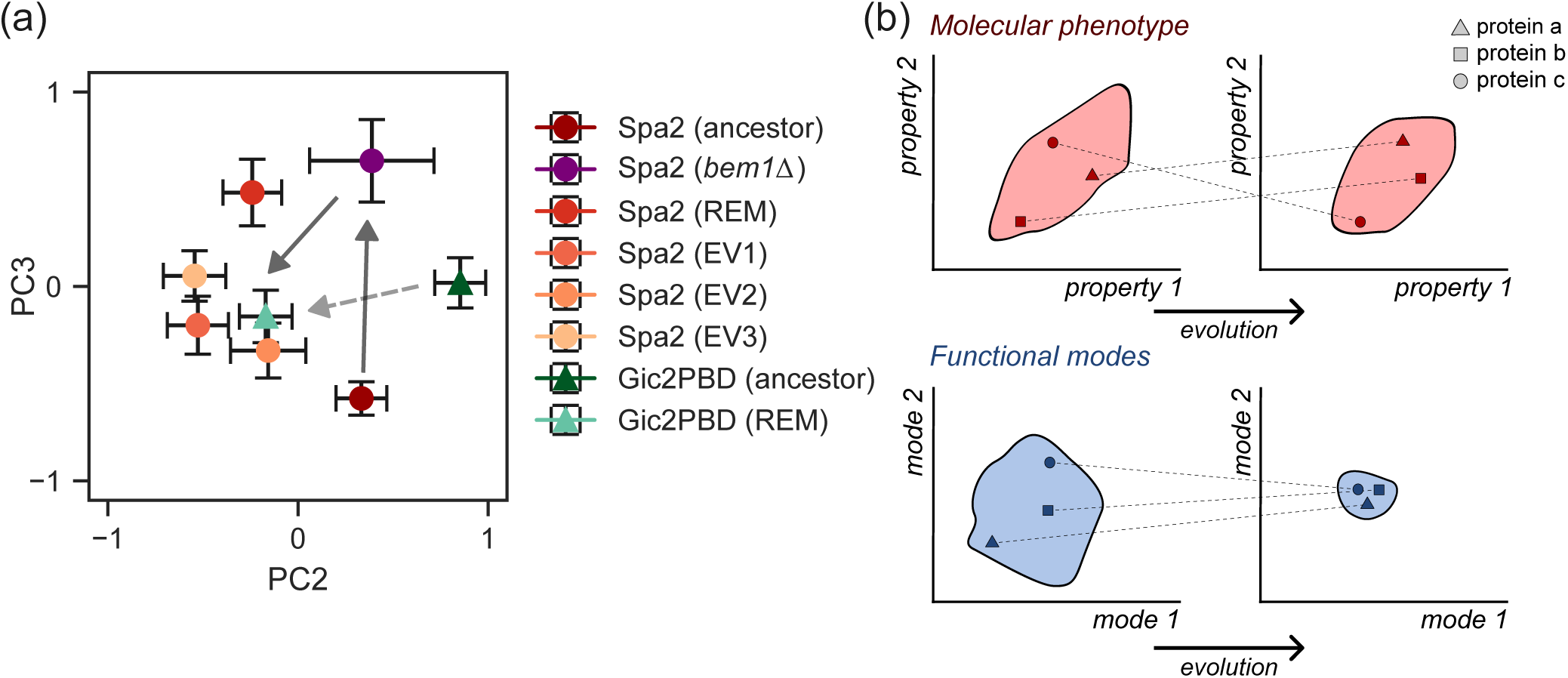
Development of PCA-extracted phenotypic modes over evolutionary recovery. (a) Means and standard errors of each marker-background combination across the phase space spanned by PC2 and PC3. REM = reconstructed evolved mutant. (b) Hypothetical conceptualization of the development of complex phenotypes as observed in the evolutionary trajectory. Individual phenotypic characteristics are variable and unconstrained across evolution, while functional behavioral modes converge to a smaller area of the available phase space.

In summary, these results reveal the presence of distinct localization modes which are largely independent of protein identity and converge across evolutionary recovery. In the context of yeast polarization, this suggests that compensatory evolution balances sufficient accumulation of proteins during polarization with global timescale restrictions, resulting in converging trends in recovered phenotypes. More broadly, these observations suggest that specific characteristics of protein dynamics, or the molecular phenotype, may not be under stringent selection; rather, evolution might shape collective behavioral modes that are bounded by functional constraints (Fig. 6(b)).

## DISCUSSION

In this work, we assessed the recovery of polarization phenotypes in terms of protein localization dynamics across evolutionary repair following a genetic perturbation. Our findings demonstrate how evolutionary recovery restored the macroscopic functionality of polarization, while significantly altering the molecular phenotype. Importantly, this effect appears to apply regardless of gene essentiality; in particular Cdc42 is a strictly essential signaling component in yeast and higher order eukaryotes^48^, yet its dynamics during polarization are not conserved in our evolutionary trajectory. We therefore expect that the changes in localization dynamics are not limited to Cdc42 and Spa2 but also affect many other proteins in the polarity network, thus establishing a collective change in response to the perturbation.

The results presented here suggest a significant level of freedom in localization patterns of individual polarity proteins. However, how this variability relates to higher-level constraints remains an open question. The variation in dynamics of successful polarization modes and the absence of strong correlative relations in phenotypic characteristics (Fig. S10(a) and (b)) indicate a lack of stringent constraints in the localization behavior of polarizing proteins. Interestingly, the development of the derived generic polarization modes, interpreted as rapid recruitment and pro-longed localization (Fig. 5(c) and (e)), suggests a shared tendency towards balancing localization timescales after loss of recruitment efficiency. This potential constraint may be functional rather than physical: Spa2 in perturbed *bem1*Δ cells occupies a more extreme position in the phase space spanned by the two polarization modes compared to the evolved mutants (Fig. 6(a)), suggesting that this area is physically accessible but not favored. It would be worthwhile to further expand the pool of evolved polarization mutants to explore potential constraints to the phase space more exhaustively.

Considering the role of gene essentiality in more detail, a recent genome-wide transposon mutagenesis screen applied to the evolutionary recovery trajectory studied here demonstrated that approximately 13% of all genes showed a significant change in disruption tolerance in the reconstructed evolved mutant compared to the wildtype ancestor^23^. Data from this study suggest that the reconstructed evolved mutant remains similarly intolerant to disruption of Cdc42 (extended data from^23^), indicating that Cdc42 did not decrease in importance in this background. Hence, it could be that presence of Cdc42-GTP, although reduced in strength, is still equally required for successful polarization. Another possibility is that Cdc42 mostly remains important in other phases of the cell cycle, for instance during cytokinesis^18,49^. By contrast, Spa2 and other polarisome members become more important in the evolved mutant^23^, seemingly in accordance with the increased presence of Spa2 at the polar zone found in this work. Another striking observation from the mutagenesis screen is that a cluster of genes associated with axial landmark recognition, such as *RSR1*, *AXL2* and *BUD2*, increased in importance in the reconstructed evolved mutant^23^, while evolved mutants show an increased probability of budding away from the axial landmark (Fig. 3(a)). It is possible that the decreased disruption tolerance of these genes in the mutagenesis screen should not be interpreted in terms of their role in bud site selection but instead points at other roles in polarization, which many of these proteins have^43,50^. More generally, these points illustrate how observations at the genetic level do not necessarily allow for straightforward predictions of protein behavior inside the cell. Additional imaging of strong hits from the transposon mutagenesis screen would be valuable to further explore the relation between gene essentiality and protein localization dynamics.

The evolutionary trajectory studied in this work is an example of recovery from a deleterious loss-of-function mutation by additional gene deletions. Evolution by gene loss is highly prevalent in evolutionary history of *S. cerevisiae* ^51,52^ as well as in other branches of the tree of life, including animals^53–55^ and plants^56,57^. In some cases, loss of genes has been shown to have an adaptive effect, for instance in response to environmental stress^58,59^. It has been debated whether defunctionalization and loss of genes is primarily a streamlining process, or if evolution by gene loss can generate phenotypic novelty^60^. Adding to existing evidence of the latter^61,62^, our results demonstrate the development of qualitatively and quantitatively different phenotypes in cells that evolved through gene-inactivating mutations. This shows how new protein dynamics can arise as a consequence of adaptive gene losses, suggesting that the molecular organization of intracellular pattern formation can be innovated by removal of components.

Overall, we would propose that our findings reflect a more general principle rather than a particular exception. Our work showed substantial differences in molecular phenotype even within mutants of the same species, separated by only a handful of mutations. However, comparative studies have revealed considerable genetic diversity in polarity networks across other fungal species^6^. Hence, it is plausible that protein dynamics in these species are not conserved either, but instead exhibit significant variation while performing the same cellular function. Similar effects have already been reported in the context of other cellular processes, such as for clathrin-mediated endocytosis^10^ or mitotic spindle assembly^4,63^. Together, these observations are indicating that many essential cellular functions have a multitude of molecular solutions, thereby allowing diversity in microscopic phenotypes while maintaining the macroscopic function. As a consequence, if cellular function is not conserved in its molecular makeup, the question becomes at which level natural selection does take place in these systems. For this, more extensive mapping of phenotypic conservation and diversification between organisms or functions is required to identify possible shared patterns. Ultimately, this may lead to a more universal understanding of complex cellular functions and their evolution.

## ACKNOWLEDGMENTS

This work was funded by the European Research Council via grant 758132. The authors thank all members of the lab for their critical feedback on the manuscript.

## METHODS

### Experimental model and study participant details

All yeast strains used in this study were derived from the *Saccharomyces cerevisiae* W303 background. All strains are listed in Table S3. Yeast were grown in rich medium (YPD: 10 g/L yeast extract, 20 g/L peptone, 20 g/L dextrose) or low fluorescence synthetic complete medium (SC-LF: 6.9 g/L nitrogen base, 790 mg/L complete amino acid mix, 20 g/L dextrose) at 30 °C in glass culture tubes placed in a rotating incubator.

### Method details

#### Yeast strain construction

Strains used in this study are listed in Table S3.

Strains yMG01b and yMG05a contain *CDC42-sfGFP*^sw^ integrated at the native locus using homologous recombination and selection with a *URA3* marker and were created as described in^24^.

Strains yLL129, yLL131 and yLL132 contain Spa2-Citrine with a GDGAGLIN linker integrated at the native locus using homologous recombination and selection with a *URA3* marker and were created as described in^22^.

Strain yLL124 was obtained by sporulating heterozygous diploid strain yLL12a originating from^22^.

Strain yMG13c was created from yLL124 by homologous recombination of 60 bp flanks upstream and downstream of *NRP1* and selection with a hphMX4 marker.

Strain yMG14a was created from yMG13c by integrating *CDC42-sfGFP*^sw^ at the native locus using homologous recombination and selection with a *URA3* marker.

Strain yMG24c contains the *GIC2* PBD domain (*GIC2* aa1-208 with mutation W23A^39,40^) under the native *GIC2* promoter (304 bp) fused to 3 copies of yeast optimized GFP (yeGFP) with a LINGTKAGGS linker between Gic2PBD and the first yeGFP and RS linkers between each copy of yeGFP integrated at the *HO* locus and was created as described in^24^.

Strain yMG25a was created the same way as yMG24c, but with ancestor yMG13c.

A1 S, A10 S and T3 S were created by transforming the final evolved populations A1, A10 and T3 from^22^ with Spa2-Citrine with a GDGAGLIN linker, integrating at the native locus using homologous recombination and selecting a single clone using a *URA3* marker.

All integrations and deletions were confirmed by PCR. In the case of integrations, modified loci were fully sequenced.

#### Growth rate assay

Cells were taken directly from glycerol stocks and suspended in 50 µl YPD. 1 µl suspension was added to 200 µl YPD in wells of a 96 well plate. Growth was monitored by measuring optical density (OD) at 600 nm using a Biotek Epoch 2 Microplate Spectrophotometer kept at 30 °C while shaking the plate. OD was measured every 7 minutes for 80+ hours. To determine the doubling time during the exponential growth phase, the log of the data was fitted to a linear relation of the form *y* = *ax* + *b* in a sliding window. The doubling time *T* was derived from the maximum slope *a* as *T* = log(2)*/a*.

#### Cell morphology assay

Cells were grown to mid-log phase (OD_600_ 0.2-0.8) in SC-LF. 1 ml culture was taken out into a sterile 1.5 ml Eppendorf tube. Cells were spun down for 5 minutes at 3000 g and 950-990 µl supernatant was removed depending on the size of the pellet. Cells were resuspended and vortexed for 1 minute before mounting 1 µl of suspension on a microscope slide. Slides were imaged on a widefield microscope (IX81 inverted microscope, Olympus) with a 40X air objective equipped with an ORCA-Flash 4.0 sCMOS camera (Hamamatsu Photonics) using the brightfield channel. Brightfield images containing *>*2000 cells per replicate were segmented by YeaZ^64^, which outputs cell mask sizes and extracts major and minor axis lengths through PCA (https://github.com/rahi-lab/YeaZ-GUI).

#### Bem1-deficient backgrounds

The study of Cdc42-sfGFP^sw^ localization in Bem1-deficient cells is based on imaging data from our previous work^24^, for which Cdc42-sfGFP^sw^ was expressed in cells where Bem1 was actively depleted using the auxin inducible degron (AID) system^65^: native Bem1 was labeled with an AID tag and cells were incubated with 0.25 mM 1-naphthaleneacetic acid (NAA) for *>*30 min before imaging, inducing degradation of Bem1^24^. Data for Spa2-Citrine was generated for this study and was obtained in true *bem1*Δ knock-out cells, obtained by sporulating a heterozygous diploid^22^.

#### Live cell imaging

Cdc42-sfGFP^sw^ and Spa2-Citrine were endogenously expressed from their native genomic loci. Gic2PBD-3xGFP was expressed under its native promoter from the HO locus. Cells were grown to mid-log phase (OD_600_ 0.2-0.8) in SC-LF. 1 ml culture was taken out into a sterile 1.5 ml Eppendorf tube. In the case of AID degradation, cells were incubated with 0.25 mM 1-naphthaleneacetic acid (Sigma-Aldrich) at 30 °C for at least 30 minutes before continuing. Cells were spun down for 5 minutes at 3000 g and 950-990 µl supernatant was removed depending on the size of the pellet. Cells were resuspended and vortexed for 1 minute before mounting 1 µl of suspension on a 35mm imaging dish with a polymer coverslip bottom (ibidi). The droplet of cells was covered by an agar pad, prepared by using a 15 ml falcon tube to cut a circular pad out of a 9 cm diameter petri dish containing 40 ml SC-LF + 3% agarose (Carl ROTH).

For all single-channel imaging, cells were imaged on a laser scanning confocal microscope (Eclipse Ti inverted microscope equipped with a A1R confocal unit, Nikon Instruments) with full environmental control (Tokai Hit) kept at 30 °C and a z-drift compensation module (Nikon Perfect Focus System). Images were captured using resonant scanning mode (unidirectional) with 4x line averaging (250 ms/frame), which was found to allow for sufficient signal levels as well as maintaining acquisition speed and cell viability. Excitation was done using a 488 nm (GFP) or 514 nm (Citrine) argon laser (Dynamic Laser) and emission filter 525/50 (GFP) or 540/30 (Citrine) through a 100X/1.49NA SR Apo TIRF oil objective (Nikon); signal was detected by an A1-DUG GaAsP Multi Detector unit with gain 50. Images (9 z-slices, 0.9 µm spacing, pinhole diameter 174 µm for GFP or 183 µm for Citrine) were taken every 20 s for a total of 3-5 hours. Images were collected from the fluorescence channel and the transmitted light detector; fluorescence images were denoised using Nikon’s Denoise.AI^66^.

For the dual-channel imaging in Fig. S8, cells were imaged on a widefield microscope (Eclipse Ti inverted microscope, Nikon Instruments) equipped with full environmental control (Okolab) kept at 30 °C and a z-drift compensation module (Nikon Perfect Focus System). Excitation was done with a CoolLED pE-4000 LED light source (GFP: 470 nm, excitation filter 470/40, emission filter 525/50; mCherry: 580 nm, excitation filter 580/25, emission filter 625/30) and images were collected through a 100X/1.45NA TIRF oil objective (Nikon) with an Andor Life 888 EM-CCD camera. Images were taken at constant z-height every 1 min for a total of 8 hours.

#### Image analysis of polarization events

The number of analyzed events per condition are given in Table S1.

Image analysis of single-channel imaging was performed as described in detail in^24^, with some minor additions. Individual yeast cells were segmented from the bleaching corrected field of view and polarization events were manually annotated from at least 5 frames before the onset of polarization until the onset of bud emergence. Polarized signal was segmented by defining a signal threshold based on the mean of the cytoplasmic signal in the unpolarized state plus a number of standard deviations (3 for Cdc42-sfGFP^sw^ and Gic2PBD-3xGFP and 4 for Spa2-Citrine). The thresholded polarized signal was then used to calculate total intensity traces and the observables describing the polarization phenotype listed in Fig. S5. Adding to the procedure from^24^, a spot separation algorithm was included to deal with events in which the polarized signal oscillates between two spots before committing to a single site (Fig. S11(a)). To separate the winning spot, polarized signal was first segmented by thresholding as described. Per frame, the center of mass of each separate 3D thresholded region was calculated. Centers of mass of all polarized regions in the full timelapse were plotted together in one 3D space and grouped into clusters using agglomerative clustering, using the average distance between sets as linkage criterion with a distance threshold of 15 (Fig. S11(b)). Parameters were optimized such that timelapses containing only a single spot were not split into different clusters. The intensity traces of spots belonging to each separate cluster were then used to identify the winning cluster, defined as the cluster with the highest total intensity in the final four frames (Fig. S11(c)). Only the winning cluster was subsequently analyzed as usual. Note that the spot separation algorithm was not applied to the analysis of dissociating spots in Bem1-deficient cells, since no winning cluster can be identified in the absence of bud emergence.

For the dual-channel imaging in Fig. S8, signal thresholds were defined per channel as the unpolarized signal mean plus a visually determined number of standard deviations. Thresholds were kept constant for each channel between ancestor and mutant.

For the budding angle distributions in Fig. 2(b), Fig. 3(a) and Fig. S3(b-d), budding angles were manually measured using the angle tool in ImageJ.

All analysis was implemented in Python unless otherwise specified.

#### Principal component analysis

Principal component analysis was performed by pooling the polarization observables from Fig. S5(f from Gic2PBD and Spa2 for all polarization events leading to successful budding, including all evolved mutants and successful events in *bem1*Δ cells in the case of Spa2. All observables were included with the exception of spot mobility during maintenance Δ*s_m_*, since the near-absence of a maintenance phase for Gic2PBD in the reconstructed evolved mutant renders this parameter meaningless for many polarization events in that background. Data were standardized by subtracting the mean and scaling to unit variance. Principal components and eigenvalues were calculated using singular value decomposition. Composition of all principal components are given in Table S2.

### Quantification and statistical analysis

The number of polarization events analyzed for each strain and/or condition are given in Table S1. Events were collected from three or more experimental replicates (i.e. independent microscopy sessions). The number of replicates for the other types of data, the meaning of plotted error bars and the applied statistical methods are mentioned in the corresponding figure legends.

