## Supplemental Information for "Diversification of budding yeast polarization dynamics after compensatory experimental evolution"

### SUPPLEMENTAL INFORMATION INDEX

Figures S1-S11 and their legends.

Tables S1-3 and their legends.

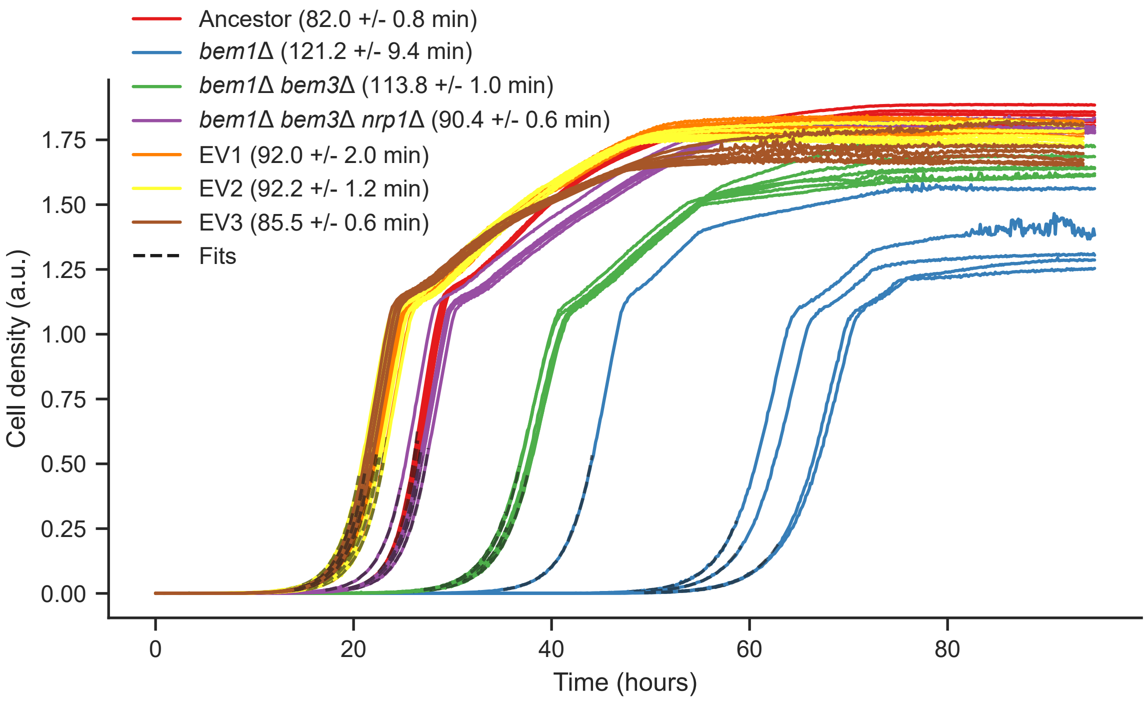


Figure S1: Population growth along the evolutionary trajectory. Growth curves of ancestral cells, *bem1*$\Delta$ knock-outs and reconstructed and evolved mutants, including intermediate mutant *bem1*$\Delta$ *bem3*$\Delta$. Numbers in the legend represent mean doubling times +/- the standard deviation. All strains express Spa2-Citrine.

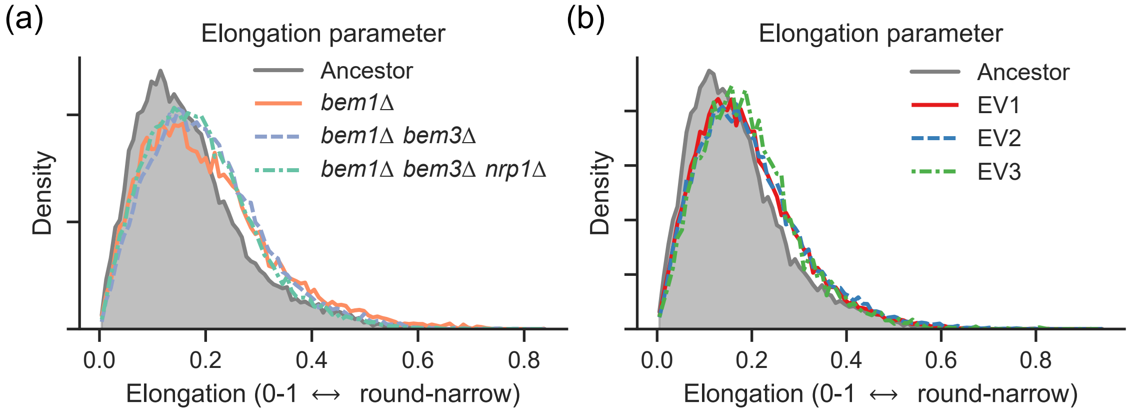


Figure S2: Cellular elongation along the evolutionary trajectory. Distributions of cell elongation $e$, defined as $e = \frac{L_{\mathrm{maj}} - L_{\mathrm{min}}}{L_{\mathrm{maj}}}$ with $L_{\mathrm{maj/min}}$ the length of the major and minor cell axis respectively. (a) Distribution of elongations along the reconstructed recovery trajectory, compared to the ancestor (grey). All strains express Spa2-Citrine. (b) Distributions of elongations of evolved mutants EV1-3 expressing Spa2-Citrine, compared to the ancestor (grey).

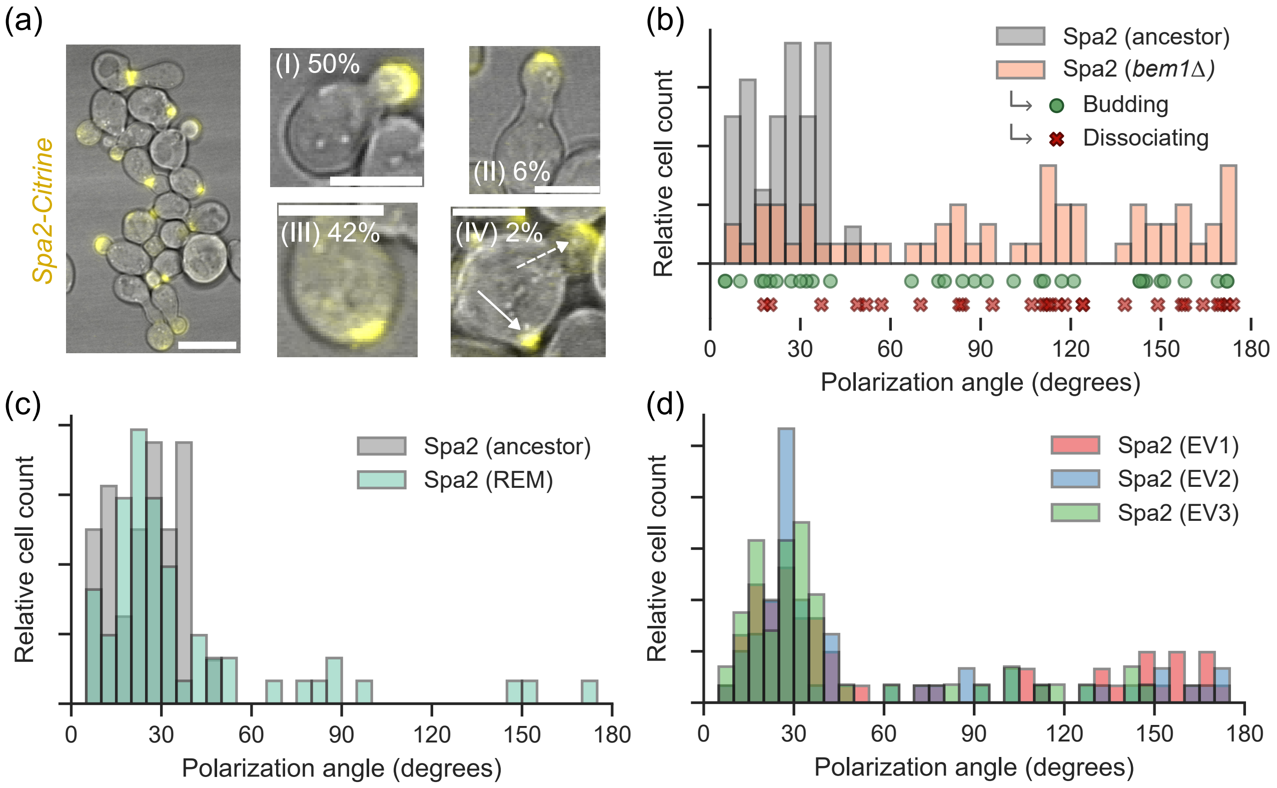


Figure S3: Characteristics of Spa2-Citrine localization in *bem1*Δ cells and in evolved mutants. (a) Representative maximum projected confocal micrographs of *bem1*Δ cells expressing Spa2-Citrine (yellow) overlaid with brightfield images. Left: large field of view (scale bar 10 µm). Right: examples of polarization phenotypes. (I) normal polarization and growth, (II), polarization resulting in elongated cell growth, (III) dissociating polarity spots and (IV) continued polarization without bud growth (solid arrow), while successfully budding elsewhere in the cell (dashed arrow). Percentages reflect phenotype occurrence in a random count of n = 100 cells. (b) Polarization angles with respect to the previous division site of Spa2-Citrine in the wildtype ancestor (grey) and *bem1*Δ cells (orange). Circles and crosses indicate whether polarization events in Bem1-depleted cells initiated successful daughter cell growth. (c) Polarization angles with respect to the previous division site of Spa2-Citrine in the wildtype ancestor (grey) and reconstructed evolved mutant (blue). (d) Polarization angles with respect to the previous division site of Spa2-Citrine in the wildtype ancestor (grey) and evolved mutants EV1 (red), EV2 (blue) and EV3 (green).

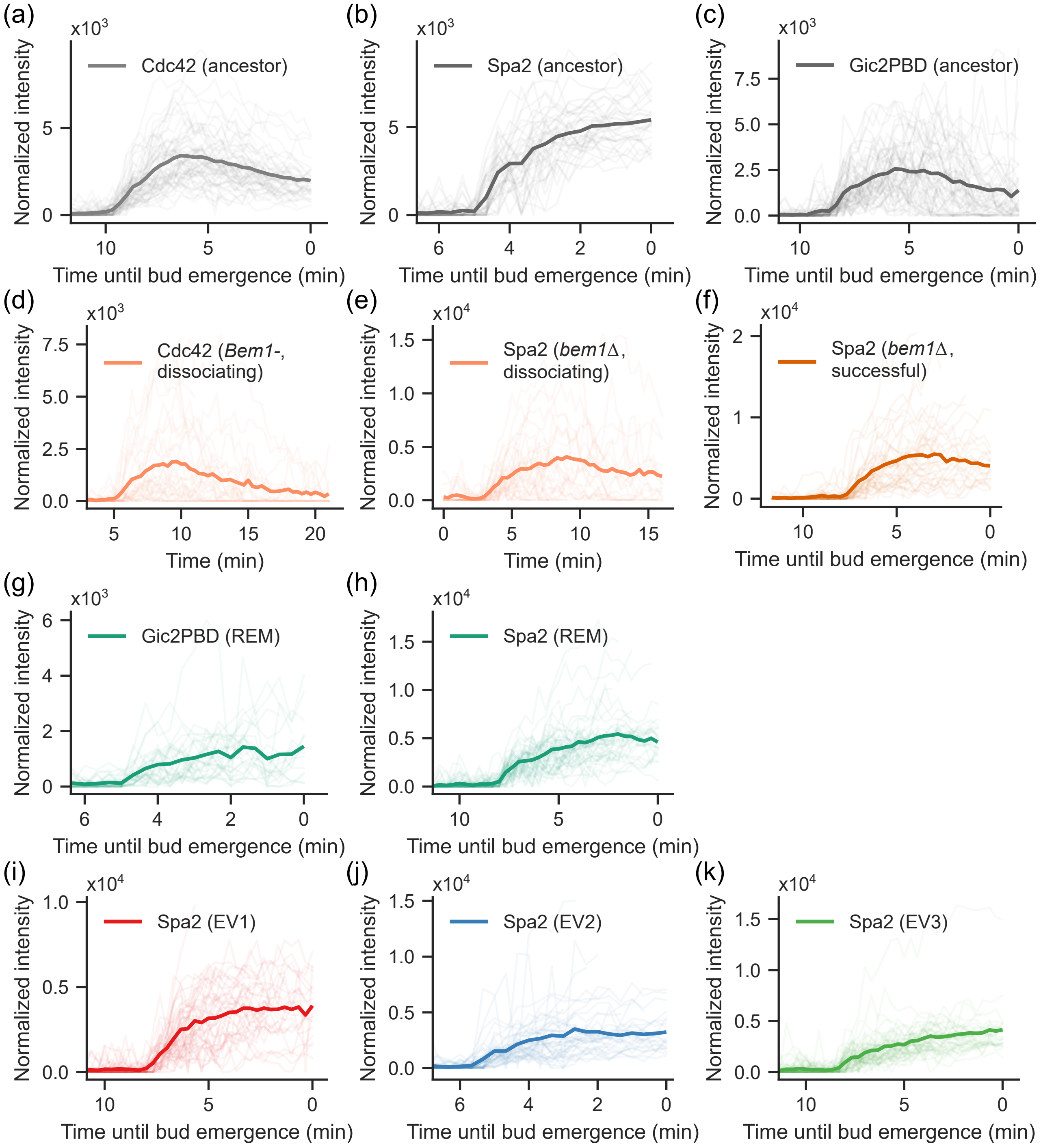


Figure S4: Individual polarized intensity traces for all markers and conditions. Bold line corresponds to the mean intensity trace as displayed in the main figures. (a) Cdc42-sfGFP$^{sw}$ in the ancestor. (b) Spa2-Citrine in the ancestor. (c) Gic2PBD-3xGFP in the ancestor. (d) Cdc42-sfGFP$^{sw}$ in Bem1-deficient cells, dissociating spots. (e) Spa2-Citrine in *bem1*Δ, dissociating spots. (f) Spa2-Citrine in *bem1*Δ, successful spots. (g) Gic2PBD-3xGFP in the reconstructed evolved mutant (REM). (h) Spa2-Citrine in the reconstructed evolved mutant (REM). (i) Spa2-Citrine in EV1. (j) Spa2-Citrine in EV2. (k) Spa2-Citrine in EV3.

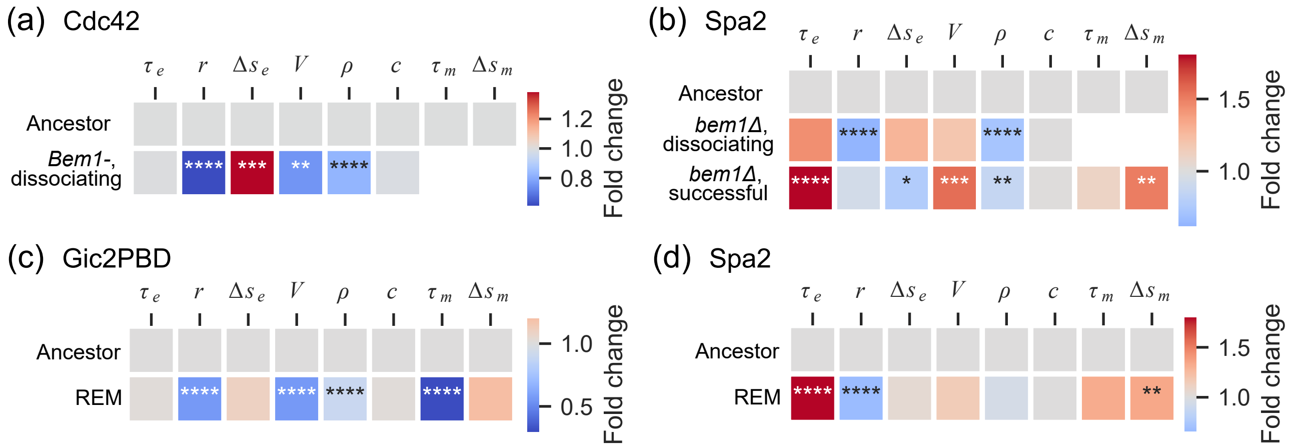


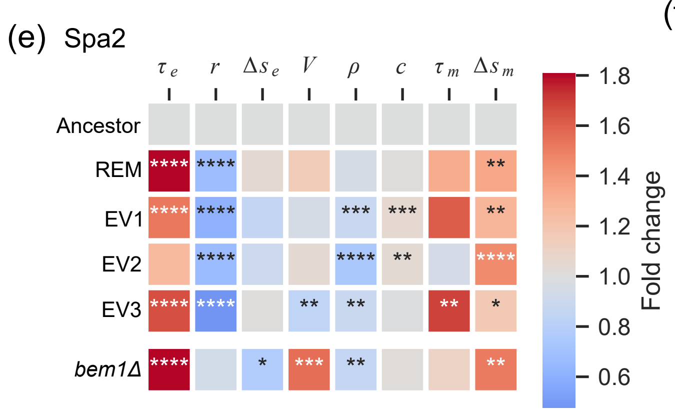


(f)

| Observable | Symbol |
|---|---|
| Establishment time | $\tau_e$ |
| Signal increase rate | $r$ |
| Mobility during establishment | $\Delta s_e$ |
| Peak spot volume | $V$ |
| Peak spot density | $\rho$ |
| Peak spot circularity | $c$ |
| Mobility during maintenance | $\Delta s_m$ |
| Maintenance time | $\tau_m$ |

Figure S5: Relative changes in polarization observables. Fold change of observable means describing the dynamics of polarizing proteins in indicated mutants with respect to the ancestor. Stars indicate significance in a Mann-Whitney U-test (*$p < 0.05$; **$p < 0.01$; ***$p < 0.001$; ****$p < 0.0001$; no symbol, not significant). (a) Cdc42-sfGFP$^{\text{sw}}$ in Bem1-deficient cells. (b) Spa2-Citrine in *bem1*$\Delta$. (c) Gic2PBD-3xGFP in the reconstructed evolved mutant (REM). (d) Spa2-Citrine in the reconstructed evolved mutant. (e) Spa2-Citrine in the reconstructed evolved mutant, evolved mutants EV1-3, and *bem1*$\Delta$ for reference. (f) Definition of observables from[24].

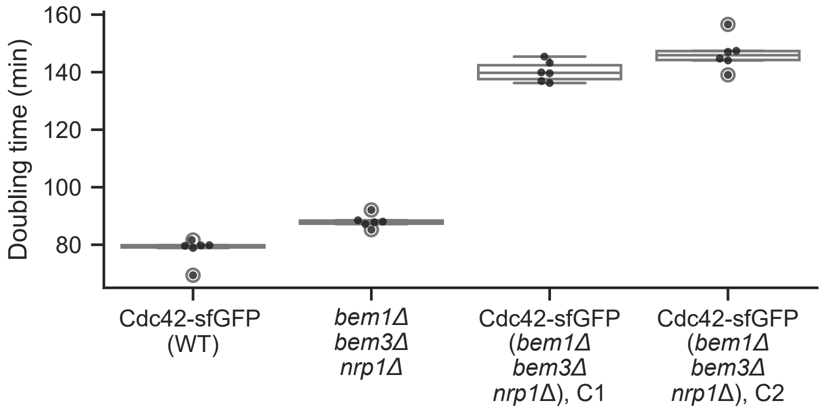


Figure S6: Growth impairment by expression of Cdc42-sfGFP$^{sw}$ in the reconstructed evolved mutant. Doubling times of cells expressing Cdc42-sfGFP$^{sw}$ in a wildtype background and in two biological replicates (C1 and C2) of integration into the *bem1*Δ *bem3*Δ *nrp1*Δ mutant. Doubling time of *bem1*Δ *bem3*Δ *nrp1*Δ cells is shown for reference.

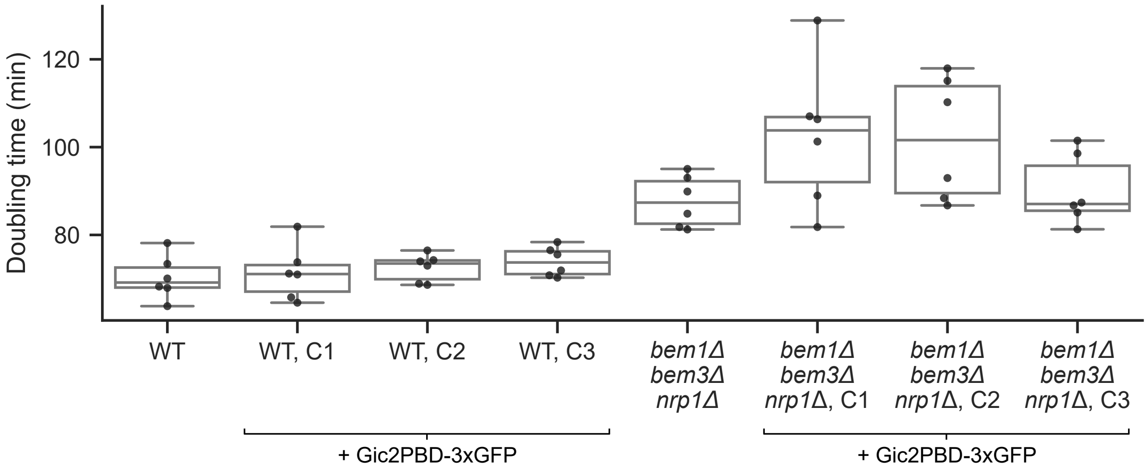


Figure S7: Effect of Gic2PBD expression on growth rate. Doubling times of cells expressing Gic2PBD-3xGFP in a wildtype background and the *bem1*Δ *bem3*Δ *nrp1*Δ mutant, three biological replicates (C1-3) each. Doubling time of strain ancestors is shown for reference.

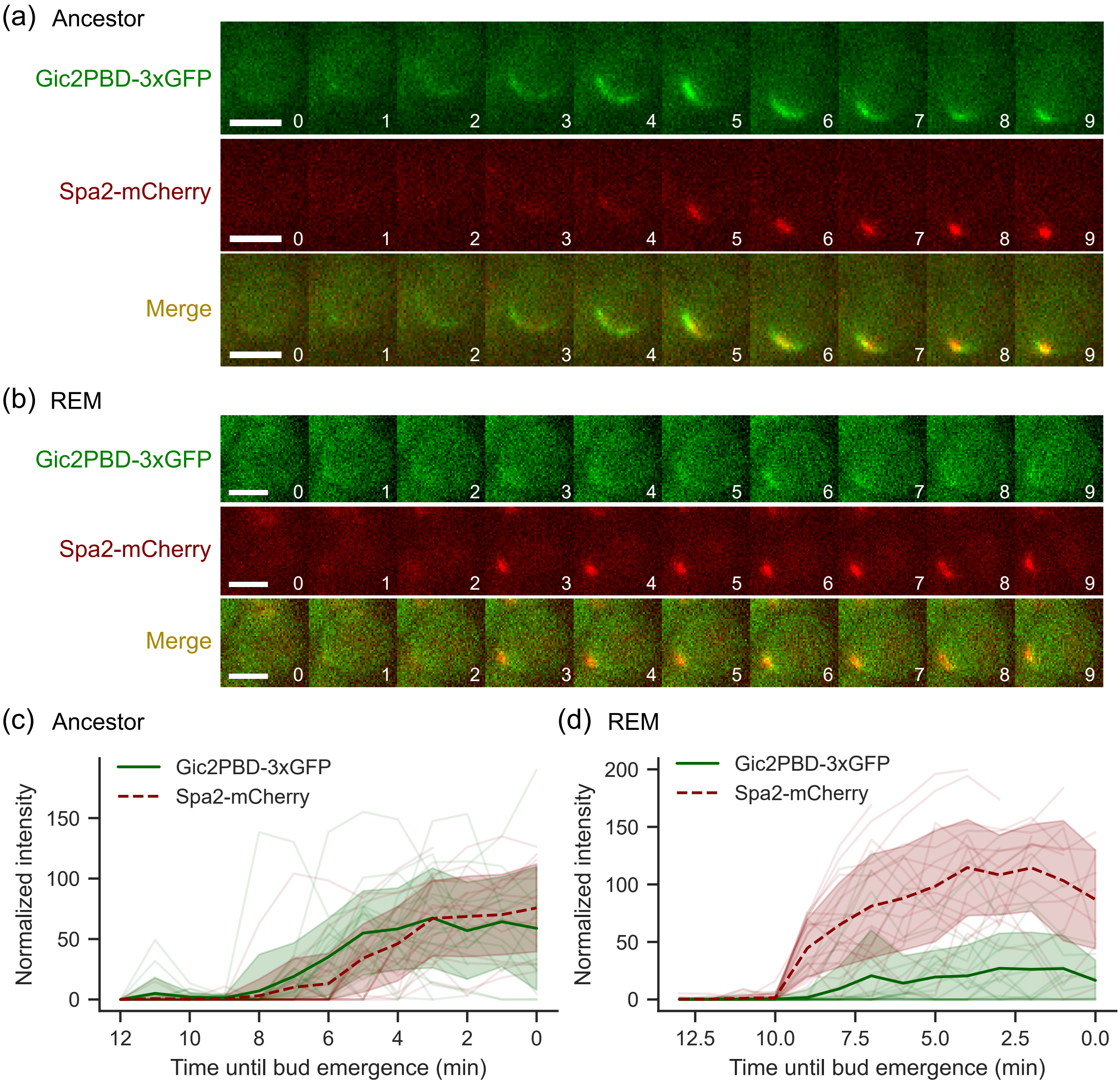


Figure S8: Relative polarization dynamics of Gic2PBD and Spa2 in the ancestor and the reconstructed evolved mutant. (a) Representative widefield fluorescence micrographs of a polarizing ancestral cell expressing Gic2PBD-3xGFP (green) and Spa2-mCherry (red). Scale bars 3 µm. Numbers indicate time in minutes. (b) Same as (a), but for a reconstructed evolved mutant (REM) cell. (c) Normalized intensity traces of polarized (thresholded) Gic2PBD-3xGFP (green) and Spa2-mCherry (red) in the ancestor (n = 20), aligned with respect to the first onset of polarization in either of the channels. (d) Same as (c), but in the reconstructed evolved mutant (REM, n = 20).

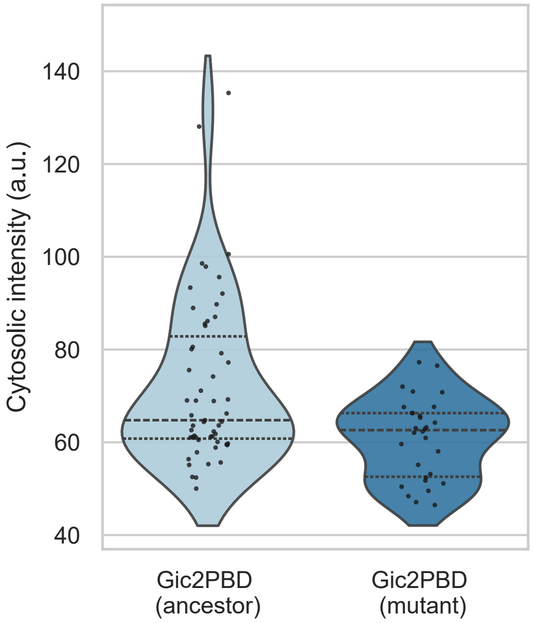


Figure S9: Expression levels of Gic2PBD-3xGFP. Green fluorescence intensity in the unpolarized cytosol of cells expressing Gic2PBD-3xGFP, obtained by taking the mean intensity of the first 5 (maximum z-projected) frames of each polarization event, in the wildtype ancestor and reconstructed evolved mutant, normalized by the laser power used in each experiment; other imaging parameters were kept identical. Dashed lines indicate data quartiles.

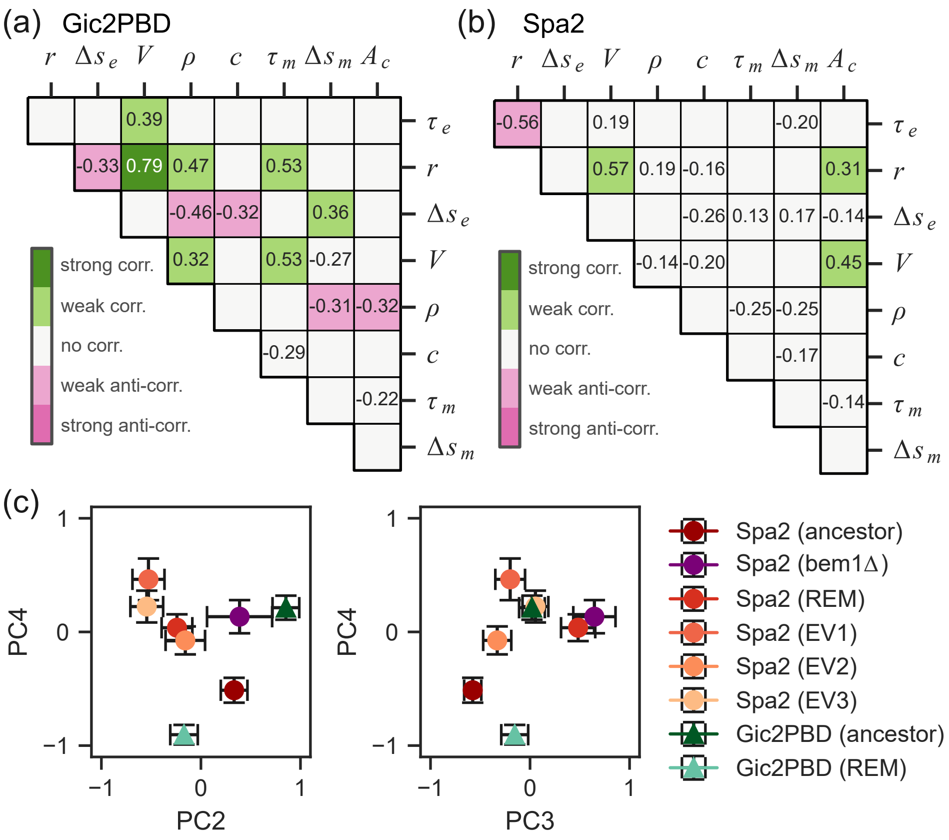


Figure S10: Dimensional reduction in polarization observable space. (a) Spearman's correlation coefficients $r_s$ of all observables as well as cell size $(A_c)$ for Gic2PBD-3xGFP in the studied backgrounds (ancestor and reconstructed evolved mutant) combined. No number is given for insignificant correlations ($p < 0.05$). Color coding indicates strong ($|r_s| > 0.6$), weak ($0.3 < |r_s| < 0.6$) or no ($|r_s| < 0.3$) (anti-)correlations. (b) Same as (a), but for Spa2-Citrine in all studied backgrounds (ancestor, successful polarization events in *bem1*Δ, reconstructed evolved mutant and EV1-3). (c) Means and standard errors of each marker-background combination across the phase space spanned by PC2-4. REM = reconstructed evolved mutant.

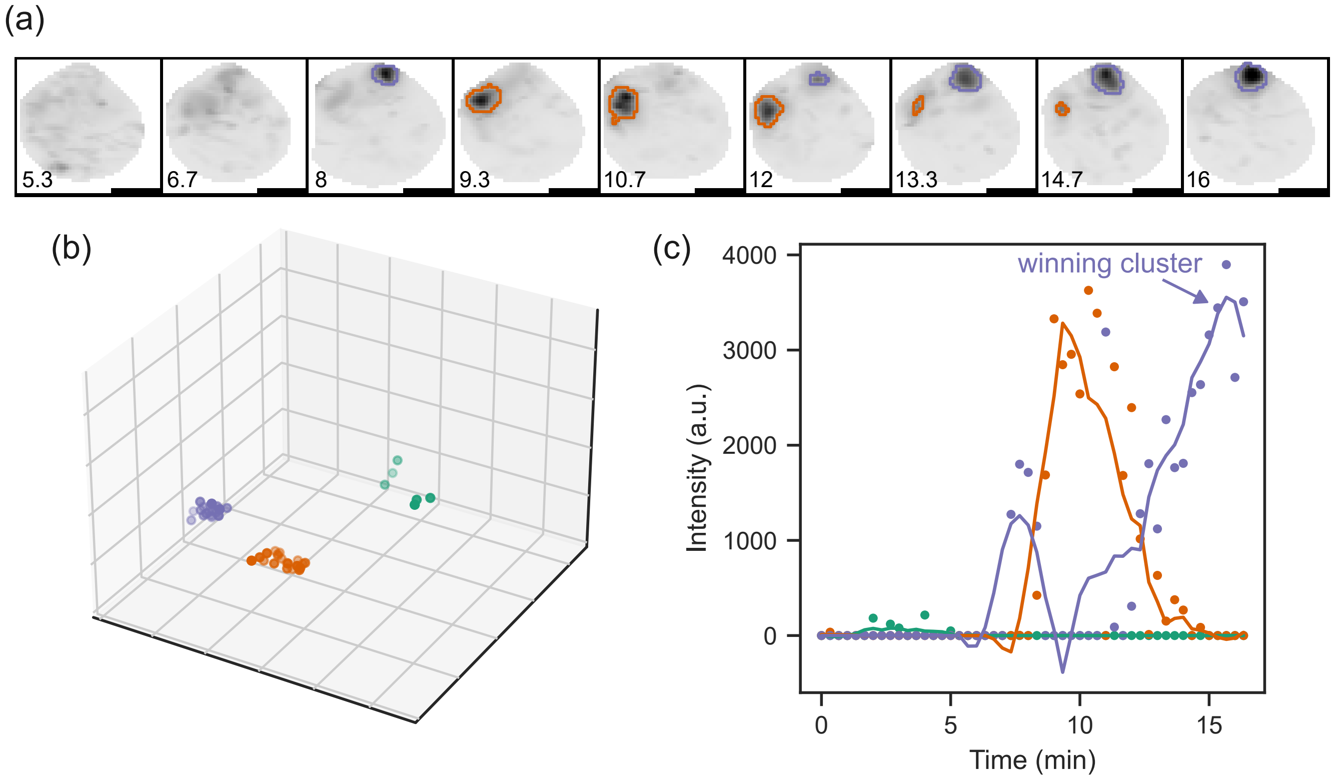


Figure S11: Spot separation algorithm. (a) Representative example of Spa2-Citrine showing oscillations over separate spots. Different colored outlines correspond to the clusters identified by the spot separation algorithm. Numbers indicate time in minutes. Scale bars 2 µm. (b) 3D plot of centers of mass of separate thresholded signal regions from all timelapse frames combined, corresponding to the polarization event from (a), grouped by agglomerative clustering. (c) Traces of thresholded intensity corresponding to each cluster as identified in (b).

| **Condition** | **# events** |
|---|---|
| Cdc42-sfGFP$^{sw}$ in ancestor | 60 |
| Cdc42-sfGFP$^{sw}$ in Bem1-depleted (dissociating) | 31 |
| Gic2PBD-3xGFP in ancestor | 55 |
| Gic2PBD-3xGFP in *bem1*Δ *bem3*Δ *nrp1*Δ | 30 |
| Spa2-Citrine in ancestor | 39 |
| Spa2-Citrine in *bem1*Δ (dissociating) | 33 |
| Spa2-Citrine in *bem1*Δ (successful) | 38 |
| Spa2-Citrine in *bem1*Δ *bem3*Δ *nrp1*Δ | 46 |
| Spa2-Citrine in EV1 | 44 |
| Spa2-Citrine in EV2 | 45 |
| Spa2-Citrine in EV3 | 52 |

Table S1: Number of analyzed polarization events per condition.

| | $\tau_e$ | $r$ | $\Delta s_e$ | $V$ | $\rho$ | $c$ | $\tau_m$ |
|---|---|---|---|---|---|---|---|
| **PC1** | 0.03 | 0.46 | -0.49 | 0.29 | 0.50 | 0.40 | -0.26 |
| **PC2** | -0.35 | 0.55 | 0.16 | 0.55 | -0.22 | -0.38 | 0.24 |
| **PC3** | 0.82 | -0.06 | 0.22 | 0.48 | 0.02 | -0.16 | -0.12 |
| **PC4** | 0.19 | -0.10 | -0.26 | 0.13 | -0.09 | 0.35 | 0.86 |
| **PC5** | 0.01 | 0.13 | 0.46 | -0.22 | 0.76 | -0.17 | 0.34 |
| **PC6** | 0.11 | -0.11 | -0.64 | -0.12 | 0.16 | -0.72 | 0.10 |
| **PC7** | 0.39 | 0.67 | -0.00 | -0.56 | -0.29 | -0.01 | 0.04 |

Table S2: Composition of PCA components in terms of original variables.

| Strain name | Main text/SI name | Relevant genotype | Mating type | Origin |
|---|---|---|---|---|
| yLL3a | Wildtype/ancestor | - | alpha | [22] |
| yLL124 | *bem1Δ bem3Δ* | *bem1::kanMX* \\ *bem3::natMX4* | a | [22] |
| yMG13c | *bem1Δ bem3Δ nrp1Δ* | *bem1::kanMX4* \\ *bem3::natMX4* \\ *nrp1::hphMX4* | a | This study |
| yMG01b | Cdc42-sfGFP$^{sw}$ in ancestor | *cdc42::CDC42-sfGFP*$^{sw}$*_URA3* | alpha | [24] |
| yMG14a | Cdc42-sfGFP$^{sw}$ in *bem1Δ bem3Δ nrp1Δ* | *cdc42::CDC42-sfGFP*$^{sw}$*_URA3* \\ *bem1::kanMX4* \\ *bem3::natMX4* \\ *nrp1::hphMX4* | a | This study |
| yLL132a | Spa2-Citrine in ancestor | *can1::Pmfa-HIS3* \\ *spa2::SPA2-Citrine_URA3* | a | [22] |
| yMG05a | Cdc42-sfGFP$^{sw}$ in *Bem1-* | *cdc42::CDC42-sfGFP*$^{sw}$*_URA3* \\ *bem1::BEM1-AID* \\ *YPRCtau3::OsTIR1-3xMyc* | alpha | [24] |
| yLL128a | Spa2-Citrine in *bem1Δ* | *can1::Pmfa-HIS3* \\ *spa2::SPA2-Citrine_URA3* \\ *bem1::kanMX* | alpha | [22] |
| yLL129a | Spa2-Citrine in *bem1Δ bem3Δ* | *can1::Pmfa-HIS3* \\ *spa2::SPA2-Citrine_URA3* \\ *bem1::kanMX* \\ *bem3::natMX4* | a | [22] |
| yLL131a | Spa2-Citrine in *bem1Δ bem3Δ nrp1Δ* | *can1::Pmfa-HIS3* \\ *spa2::SPA2-Citrine_URA3* \\ *bem1::kanMX4* \\ *bem3::natMX4* \\ *nrp1::hphMX4* | a | [22] |
| yMG24c | Gic2PBD-3xGFP in ancestor | *HO::GIC2PBD-3yeGFP_URA3* | alpha | [24] |
| yMG25a | Gic2PBD-3xGFP in *bem1Δ bem3Δ nrp1Δ* | *HO::GIC2PBD-3yeGFP_URA3* \\ *bem1::kanMX4* \\ *bem3::natMX4* \\ *nrp1::hphMX4* | a | This study |
| A1_S | Evolved line EV1 | *can1::Pmfa-HIS3* \\ *spa2::SPA2-Citrine_URA3* \\ *bem1::kanMX4* | a | [22] |
| A10_S | Evolved line EV2 | *can1::Pmfa-HIS3* \\ *spa2::SPA2-Citrine_URA3* \\ *bem1::kanMX4* | a | [22] |
| T3_S | Evolved line EV3 | *can1::Pmfa-HIS3* \\ *spa2::SPA2-Citrine_URA3* \\ *bem1::kanMX4* | a | [22] |
| yMG31a | Gic2PBD-3xGFP & Spa2-mCherry in ancestor | *HO::GIC2PBD-3yeGFP_URA3* \\ *spa2::Spa2-mCherry_HIS3* | alpha | This study |
| yMG32a | Gic2PBD-3xGFP & Spa2-mCherry in *bem1Δ bem3Δ nrp1Δ* | *HO::GIC2PBD-3yeGFP_URA3* \\ *spa2::Spa2-mCherry_HIS3* \\ *bem1::kanMX4* \\ *bem3::natMX4* \\ *nrp1::hphMX4* | a | This study |

Table S3: Yeast strains used in this study. All strains are haploid in a W303 background. All strains have *BUD4* replaced by *BUD4* from S288c and *leu2-3*, *his3-11*, *trp1-1* and *ura3Δ*.
